# Whole-brain modeling of the differential influences of Amyloid-Beta and Tau in Alzheimer’s Disease

**DOI:** 10.1101/2022.10.30.514365

**Authors:** Gustavo Patow, Leon Stefanovski, Petra Ritter, Gustavo Deco, Xenia Kobeleva, the Alzheimer’s Disease Neuroimaging Initiative

**Affiliations:** ViRVIG, Universitat de Girona, Girona, Spain; Berlin Institute of Health at Charité, Universitätsmedizin Berlin, Berlin, Germany; Department of Neurology with Experimental Neurology, Brain Simulation Section, Charité – Universitätsmedizin Berlin, corporate member of Freie Universität Berlin and Humboldt-Universität zu Berlin, Berlin, Germany; Center for Brain and Cognition, Computational Neuroscience Group, Department of Information and Communication Technologies, Universitat Pompeu Fabra, Barcelona, Spain; Clinic for Neurology, University Hospital Bonn, Bonn, Germany; German Center for Neurodegenerative Diseases (DZNE), Bonn, Germany; Department of Science, University of Cambridge, London, UK

**Keywords:** Alzheimer’s disease, Amyloid-Beta, Tau, Whole-Brain model, Simulation

## Abstract

**Background:** Alzheimer’s Disease is a neurodegenerative condition associated with the accumulation of two misfolded proteins, amyloid-beta (A*β*) and tau. We study their effect on neuronal activity, with the aim of assessing their individual and combined impact.

**Methods:** We use a whole-brain dynamic model to find the optimal parameters that best describe the effects of A*β* and tau on the excitation-inhibition balance of the local nodes.

**Results:** We found a clear dominance of A*β* over tau in the early disease stages (MCI), while tau dominates over A*β* in the latest stages (AD). We identify crucial roles for A*β* and tau in complex neuronal dynamics and demonstrate the viability of using regional distributions to define models of large-scale brain function in AD.

**Conclusions:** Our study provides further insight into the dynamics and complex interplay between these two proteins, opening the path for further investigations on biomarkers and candidate therapeutic targets in-silico.

## 1 Background

Alzheimer’s Disease (AD) is a neurodegenerative disease that leads to progressive impairment of memory and other cognitive domains, neuropsychiatric symptoms, and, ultimately, severe impairment in all body functions. This results in both a huge loss of quality of life for affected people and caregivers and high costs for society at large. AD pathogenesis is associated with several interlinked pathomechanistic processes, from genomics to connectomics, including the Notch-1 pathway, neurotransmitters, polygenetic factors, neuroinflammation, and neuroplasticity [1]. However, the accumulation of misfolded proteins is considered the pathological hallmark of AD: namely extracellular accumulation of Amyloid-beta (A*β*), forming senile plaques; and intraneuronal aggregation of the microtubule protein tau, called neurofibrillary tangles [2]. Treatments for removal of A*β* (e.g., with Adacanumab and Lecanemab) are currently discussed in light of inconclusive effects on reducing cognitive decline [3]. In spite of the large body of research on AD, many aspects regarding pathophysiology and the roles of A*β* and tau are still incompletely understood [4, 5].

Regarding brain dysfunction, several human autopsy and animal studies have seen a disruption in excitation/inhibition (E/I) balance, especially in early stages where neuronal hyperexcitability impairs cortical activity and thus contributes to cognitive decline [6, 7]. Chang et al. [8] showed tau affects excitatory and inhibitory neurons differently, and its removal decreases the baseline activity of excitatory neurons and, simultaneously, affects the axon initial segments and the intrinsic excitability of inhibitory neurons, resulting in network inhibition. In this line, Bi and co-workers [9] hypothesized that A*β* impairs GABAergic function and thus produces synaptic hyperexcitation. Petrache et al. [10] found synaptic hyperexcitation and severely disrupted E/I inputs onto principal cells, and a reduction in the somatic inhibitory axon terminals. Recently, Lauterborn and coauthors [11] revealed significantly elevated E/I ratios in post-mortem cortex samples. While interesting results regarding E/I imbalance with marked hyperexcitability were derived in animals and post-mortem human cortex samples, in-vivo human studies are lacking, as the activity of E/I populations cannot be directly measured using neuroimaging. Most works on whole-brain dynamics studied activation patterns but were not informative regarding the role of E/I populations [12, 13, 14, 15, 16]. To understand the complex interplay between pathophysiological processes and brain activity (i.e., fMRI), models might improve when incorporating heterogeneity of brain dynamics based on empirical data [17, 18, 19].

Earlier work using whole-brain simulations focused on linking global and local brain dynamics to individual differences in cognitive performance scores from different conditions [12]. Demirtaş [14] et al. studied the effect of heterogeneity of local synaptic strengths on a dynamical model of human cortex in healthy subjects, showing that heterogeneity significantly improved the fitting of resting-state functional connectivity. Stefanovski and co-authors [15] focused on the connection of A*β* with neural function in The Virtual Brain [20] to examine how A*β* modulates regional E/I balance, producing local hyperexcitation in regions with high A*β* loads. This led to further improvements on classifications between AD and controls [16]. However, all these works study the effect of a single burden, namely A*β*, on the neuronal dynamics, while our work focuses mostly on the *interplay* of both burdens, i.e., A*β* and tau, assessing their relative impacts on brain dynamics.

In this paper, we use whole-brain modeling techniques to study the impact of both A*β* and tau on the dynamics of regional behaviors in AD, discerning the impact of each protein in isolation and in combination, and being able to assess their relative weights on contributing to abnormal brain activity. We use the Balanced Excitation-Inhibition (BEI) model [18], which can reproduce the fMRI activity based on interactions of excitatory and inhibitory neural populations interconnected by white matter tracts. We show in this work a clear dominance of the effects of A*β* over tau on brain dynamics in the earlier stages of the disease (Mild Cognitive Impairment, MCI), and a dominance of protein tau over A*β* in advanced stages (manifest dementia).

## 2 Methods

### 2.1 Methods Overview

#### Model Creation

Figure 1a presents an overview of our overall analysis strategy, and the details could be found in the Methods Section. We make use of MRI and positron emission tomography (PET) from the Alzheimer’s Disease Neuroimaging Initiative (ADNI). In summary, we use diffusion MRI to generate the structural connectomes of healthy controls (HC), mild cognitive impairment (MCI) and Alzheimer’s Disease (AD) subjects. We use task-free resting-state functional MRI to fit a whole-brain model in which the local neuronal dynamics of each brain region evolves according to the dynamic mean field model by Deco et al. [18], which is then connected to a spontaneous blood-oxygenation-level-dependent (BOLD) dynamics. We refer to this model as the *Balanced Excitation-Inhibition (BEI)* model, which can be thought of as a homogeneous reference against which we evaluate the performance of our heterogeneous AD model. A*β* and tau distributions are derived from AV-45 and AV-1451 PET from ADNI. For the heterogeneous model, we incorporate regional heterogeneous distributions of the main proteins involved in AD, namely A*β* and tau, as first order multiplicative polynomials for each burden and for each type of population (excitatory/inhibitory) into the local gain parameter *M*_(*E,I*)_. Fitting the model to empirical fMRI data allows us to evaluate which effect of A*β* and tau to the different populations can mechanistically explain the observed behaviors.

**Figure 1.**
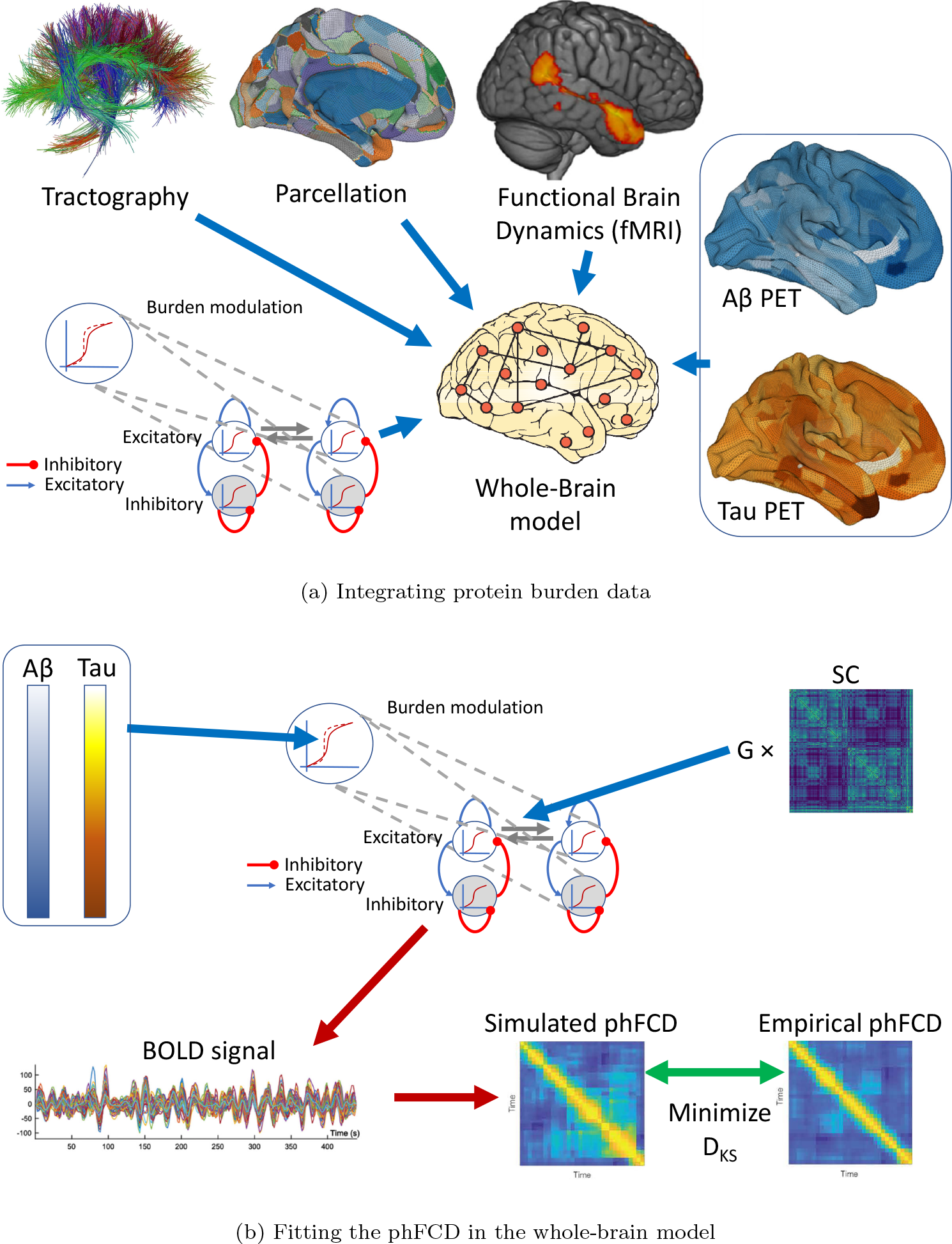
Illustrative overview of our processing pipeline. ((a)) Basic ingredients for the integration of protein burden data from structural (dMRI, top left), functional (fMRI, top right), and burden (PET, right) using the same parcellation for each neuroimaging modality (top, middle) for generating a whole-brain computational model (bottom left). Each node of the model is using a realistic underlying biophysical neuronal model including AMPA (blue connections), GABA (red), and NMDA (gray) synapses as well as neurotransmitter gain modulation of these. ((b)) Fitting the measures in the whole-brain model: First, we simulate the BOLD timeseries for each brain region in the parcellation, for each subject. These timeseries are defined by its inputs, namely a previously found global coupling constant *G*, an individual Structural Connectivity (*SC*) matrix, and the corresponding individual A*β* and tau burdens. Subsequently, we compute a time-versus-time matrix of phase functional connectivity dynamics (phFCD). This is compared to a reference empirical phFCD extracted from the fMRI data off the same subject using the Kolmogorov-Smirnov distance (KS), *D*_*KS*_, which is minimized to find the generative parameters of the model. This process is repeated for the other two measures of brain dynamics, functional connectivity (FC) and sliding-window functional connectivity dynamics (swFCD).

#### Model Fitting

For both of our models, homogeneous and heterogeneous, we assume that all diffusion MRI-reconstructed streamline fibers have the same conductivity and thus the coupling between different brain areas is scaled by a single global parameter, *G*. We first tune the *G* parameter of the BEI model to adjust the strength of effective coupling in the model and identify the brain’s dynamic working-point by fitting the model to three empirical properties that are estimated from the empirical fMRI data:

- the Pearson correlation between model and empirical estimates of static (i.e., time-averaged) functional connectivity estimated across all pairs of brain regions (FC);
- similarity in sliding-window functional connectivity dynamics (swFCD);
- the KS distance between a set of dynamic functional connectivity matrices (also called coherence connectivity matrix [21]) built from the average BOLD time series of each ROI, which were Hilbert-transformed to yield the phase evolution of the regional signals (phFCD).

We then fit the coefficients for the two burdens, for excitatory and inhibitory populations, with a global optimization algorithm, within directional bounds obtained from previous clinical observations (see below, in Section 2.8).

#### Result Evaluation

To demonstrate that E/I imbalance is dependent on the precise distribution of the A*β* and tau burdens, at the optimal values obtained with the fitting procedure described above, we randomly shuffled the empirical protein burdens; i.e., the original 378 values for each of the misfolded protein maps were randomly re-assigned to different regions, and the model was run 10 times with each different randomly re-assigned receptor map, and the simulation was repeated 10 more times for each re-assigned receptor map, for a total of 100 simulations each time. Figure 1b shows the results of randomly shuffling the empirical burden densities across the regions at the optimum point. This randomly reshuffled manipulation yields a significantly worse fit compared to the actual empirical burden densities (as shown by the Wilcoxon statistics in the boxplot). We additionally evaluate the quality of the simulation results with the optimized parameters with original (i.e., not shuffled) burdens and with the homogeneous BEI model. Finally, we examine the relevance of each type of burden by optimizing them in isolation from each other (i.e., zeroing the other one out), and comparing the results. The whole comparisons include both burdens in isolation, both burdens simultaneously, and with the homogeneous (i.e., BEI) model.

### 2.2 Participants

Empirical data were obtained from the Alzheimer’s Disease Neuroimaging Initiative (ADNI) database (`adni.loni.usc.edu`), which is a longitudinal multi-site study designed to develop biomarkers for Alzheimer’s disease (AD) across all stages. The inclusion criteria for AD patients was the NINCDS-ADRDA criteria, which contains only clinical features [22], and an MMSE score below 24. For both HC and MCI, the inclusion criteria were a MMSE (Mini Mental State Examination) score between 24-30, as well as age between 55-90 years. Also, for MCI, participants had to have a subjective memory complaint and abnormal results in another neuropsychological memory test. Imaging and biomarkers were not used for the diagnosis.

### 2.3 Data Acquisition and Processing

All the data in this study were previously used in Stefanovski et al. [15] work, so we will present here an abridged version of the processing performed on the original data and refer to the original work for the details. All images used in this study were taken from ADNI-3, using data from Siemens scanners with a magnetic field strength of 3T.

#### 2.3.1 Structural MRI

For each included participant, we created a brain parcellation for our structural data using FLAIR, following the minimal preprocessing pipeline [23] of the Human Connectome Project (HCP) using Freesurfer^[1]^ [24], FSL [25, 26, 27] and connectome workbench^[2]^. Therefore, we used T1 MPRAGE, FLAIR and fieldmaps for the anatomical parcellation. We then registered the subject cortical surfaces to the parcellation of Glasser et al. [28] using the multimodal surface matching (MSM) tool [29]. In this parcellation, there were 379 regions: 180 left and 180 right cortical regions, 9 left and 9 right subcortical regions, and 1 brainstem region.

#### 2.3.2 PET Images

For A*β*, we used the version of AV-45 PET already preprocessed by ADNI, using a standard image with a resolution of 1.5mm cubic voxels and matrix size of 160 *×* 160 *×* 96, normalized so that the average voxel intensity was 1 and smoothed out using a scanner-specific filter function. Then, a brainmask was generated from the structural preprocessing pipeline (HCP) and used to mask the PET image. On the other hand, to obtain the local burden of A*β*, we computed the relative intensity to the cerebellum. We received in each voxel a relative A*β* burden which is aggregated according to the parcellation used for our modeling approach. Subcortical region PET loads were defined as the average SUVR in subcortical gray matter (GM). With the help of the connectome workbench tool, using the pial and white matter surfaces as ribbon constraints, we mapped the Cortical GM PET intensities onto individual cortical surfaces. Finally, using the multimodal Glasser parcellation we derived average regional PET loads.

For tau, we also used ADNI’s preprocessed version of AV-1451 (Flortaucipir) following the same acquisition and processing, resulting in a single relative tau value for each voxel. Then, these values were also aggregated to the selected parcellation, also following the already mentioned steps. The final average regional tau loads were obtained in the Glasser parcellation.

#### 2.3.3 DWI

Individual tractographies were computed only for included HC participants, and they were averaged to a standard brain template (see below). Preprocessing was mainly done with the MRtrix3 software package^[3]^.

In particular, the following steps were performed: First, we denoised the DWI data [30], followed by motion and eddy current correction^[4]^. Then, B1 field inhomogeneity correction (ANTS N4), followed by a brainmask estimation from the DWI images. Next, we normalized the DWI intensity for the group of participants, which was used to generate a WM response function [31], and created an average response function from all participants. Next, we estimated the fiber orientation distribution and the average response function [32] using the subject normalized DWI image, to finally generate a five tissue type image. Finally, we used the iFOD2 algorithm [33] and the SIFT2 algorithm [34] to get the weighted anatomical constrained tractography [35], to end up merging all information into the Glasser connectome, resulting in a structural connectome (SC).

It is important to note that the multi-center nature of ADNI data can be problematic, with inter-site differences in acquisition, scanner and protocol, being DWI is particularly susceptible to multi-center related issues and problematic harmonization. To prevent these problems, we actually restricted the data set to just one scanner type, from Siemens. The details of the scanner metadata including the acquisition centers and used scanners are also provided in the supplementary material.

#### 2.3.4 fMRI

With respect to the processing of the fMRI data, the images were initially preprocessed in FSL FEAT and independent component analysis–based denoising (FSLFIX) following a basic pipeline [15]. Time courses for noise-labeled components, along with 24 head motion parameters, were then removed from the voxel-wise fMRI time series using ordinary least squares regression.

The resulting denoised functional data were spatially normalized to the MNI space using Advanced Normalization Tools (version 2.2.0). Mean time series for each parcellated region were then extracted, and interregional FC matrices were estimated using Pearson correlations between each pair of regional time series. Dynamic FC matrices were also calculated for the empirical data, as outlined below.

### 2.4 Generation of a Standard Brain Template

As previously done [15], we average the SCs of all HC participants, using an arithmetic mean

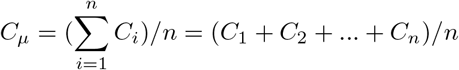

wherein *C*_*µ*_ is the averaged SC matrix, *n* is the number of HC participants and *C*_*i*_ is the individual SC matrix.

However, as matrices in this context are large (i.e., 379 regions), the average input to any given node can be too large for the DMF, making fitting and processing in general more difficult. Thus, we discarded the traditional normalization of dividing the matrix elements by its maximum, and used a slightly different approach, instead. First, we added one and applied the logarithm to every entry, as *lC* = *log*(*C*_*µ*_ + 1). Then, we computed the maximum input any node could receive for a unitary unit input current, *maxNodeInput* = *max*_*j*_(Σ_*i*_(*lC*_*i,j*_)), and finally we normalized by 0.7 *∗ lC/maxNodeInput*, where 0.7 was chosen to be a convenient normalization value. Observe that this constant is actually multiplying another constant *G* in the model which we fit to empirical data, so its actual value can safely be changed.

In Figure 3 we can find the SC matrix and organization graph, where we can observe that the general characteristics of physiological SCs such as symmetry, laterality, homology, and subcortical hubs are maintained in the averaged connectome. The election of the averaged SC allowed us to control all factors (e.g., atrophy), which matched our objective of simulating the activity from both healthy and “pathogenic” modifications by A*β* and tau.

### 2.5 Balanced Excitation-Inhibition (BEI) model

In this work we used the Dynamic Mean Field (DMF) model proposed by Deco et al. [18], which consists of a network model to simulate spontaneous brain activity at the whole-brain level. Following the original formulation, each node represents a region of interest (i.e., a brain area) and the links represent the white matter connections between them. In turn, each node is a reduced representation of large ensembles of interconnected excitatory and inhibitory integrate-and-fire spiking neurons (as in the original, respectively 80% and 20% neurons), to a set of dynamical equations describing the activity of coupled excitatory (*E*) and inhibitory (*I*) pools of neurons, based on the original reduction of Wong and Wang [36]. In the DMF model, excitatory synaptic currents, *I*(*E*), are mediated by NMDA receptors, while inhibitory currents, *I*(*I*), are mediated by *GABA*_*A*_ receptors. Both neuronal pools are reciprocally connected, and the inter-area interactions occur at the excitatory level only, scaled by the structural connectivity *C*_*kj*_ (see Section 2.3.1).

To be more specific, the DMF model is expressed by the following system of coupled differential equations:

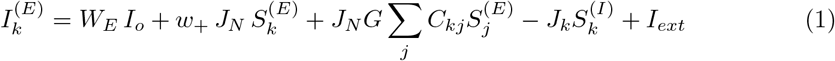

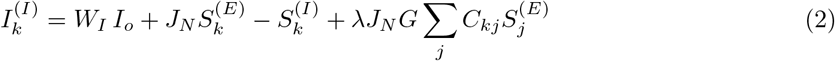

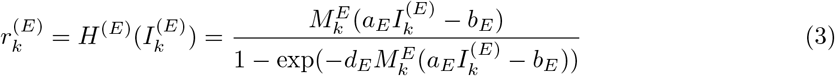

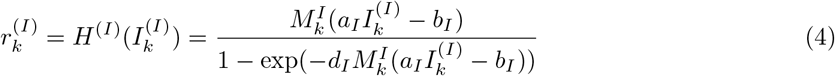

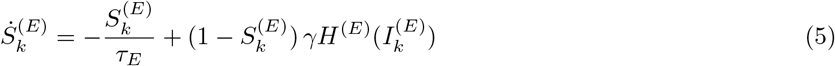

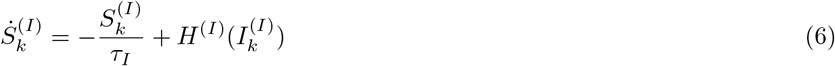

Here, the last *two* equations should add, when integrating, an uncorrelated standard Gaussian noise term with an amplitude of *σ* = 0.01*nA* (using Euler-Maruyama integration). In these equations, *λ* is a parameter that can be equal to 1 or 0, indicating whether long-range feedforward inhibition is considered (*λ* = 1) or not (*λ* = 0). In the above equation, the kinetic parameters are *γ* = 0.641*/*1000 (the factor 1000 is for expressing everything in ms), and *τ*_*E*_ = *τ*_*NMDA*_ and *τ*_*I*_ = *τ*_*GABA*_. The excitatory synaptic coupling *J*_*NMDA*_ = 0.15 (nA). The overall effective external input is *I*_0_ = 0.382 (nA) scaled by *W*_*E*_ and *W*_*I*_, for the excitatory pools and the inhibitory pools, respectively. The effective time constant of NMDA is *τ*_*NMDA*_ = 100 ms [36]. The values of *W*_*I*_, *I*_0_, and *J*_*NMDA*_ were chosen to obtain a low level of spontaneous activity for the isolated local area model. The values of the gating variables can be found in Table 1.

**Table 1.** Gating variables in the BEI model.

| Excitatory gating variables | Inhibitory gating variables |
| --- | --- |
| $a_E = 310$ ($nC^{-1}$) | $a_I = 615$ ($nC^{-1}$) |
| $b_E = 125$ (Hz) | $b_I = 177$ (Hz) |
| $d_E = 0.16$ (s) | $d_I = 0.087$ (s) |
| $\tau_E = \tau_{NMDA} = 100$ (ms) | $\tau_I = \tau_{GABA} = 10$ (ms) |
| $W_E = 1$ | $W_I = 0.7$ |

As mentioned, the DMF model is derived from the original Wong and Wang model [36] to emulate resting-state conditions, such that each isolated node displays the typical noisy spontaneous activity with low firing rate (*H*^(*E*)^ *∼* 3*Hz*) observed in electrophysiology experiments, reusing most of the parameter values defined there. We also implemented the Feedback Inhibition Control (FIC) mechanism described by Deco et al. [18], where the inhibition weight, *J*_*n*_, was adjusted separately for each node *n* such that the firing rate of the excitatory pools *H*^(*E*)^ remains clamped at 3Hz even when receiving excitatory input from connected areas. Deco et al. [18] demonstrated that this mechanism leads to a better prediction of the resting-state FC and to a more realistic evoked activity. We refer to this model as the balanced excitation-inhibition (BEI) model. Although the local adjustments in this model introduce some degree of regional heterogeneity, the firing rates are constrained to be uniform across regions so we consider this BEI model as a homogeneous benchmark against which we evaluate more sophisticated models that allow A*β* and tau to affect intrinsic dynamical properties across regions.

Following the Glasser parcellation [23], we considered *N* = 379 brain areas in our whole-brain network model. Each area *n* receives excitatory input from all structurally connected areas into its excitatory pool, weighted by the connectivity matrix, obtained from dMRI (see Section 2.3.3). Furthermore, all inter-area E-to-E connections are equally scaled by a global coupling factor *G*. This global scaling factor is the only control parameter that is adjusted to move the system to its optimal working point, where the simulated activity maximally fits the empirical resting-state activity of healthy control participants. Simulations were run for a range of *G* between 0 and 5.5 with an increment of 0.05 and with a time step of 1 ms. For each *G*, we ran 200 simulations of 435 s each, in order to emulate the empirical resting-state scans from 17 participants. The optimum value found, for the *phFCD* observable, is for *G* = 3.1. See Figure 2A.

**Figure 2.**
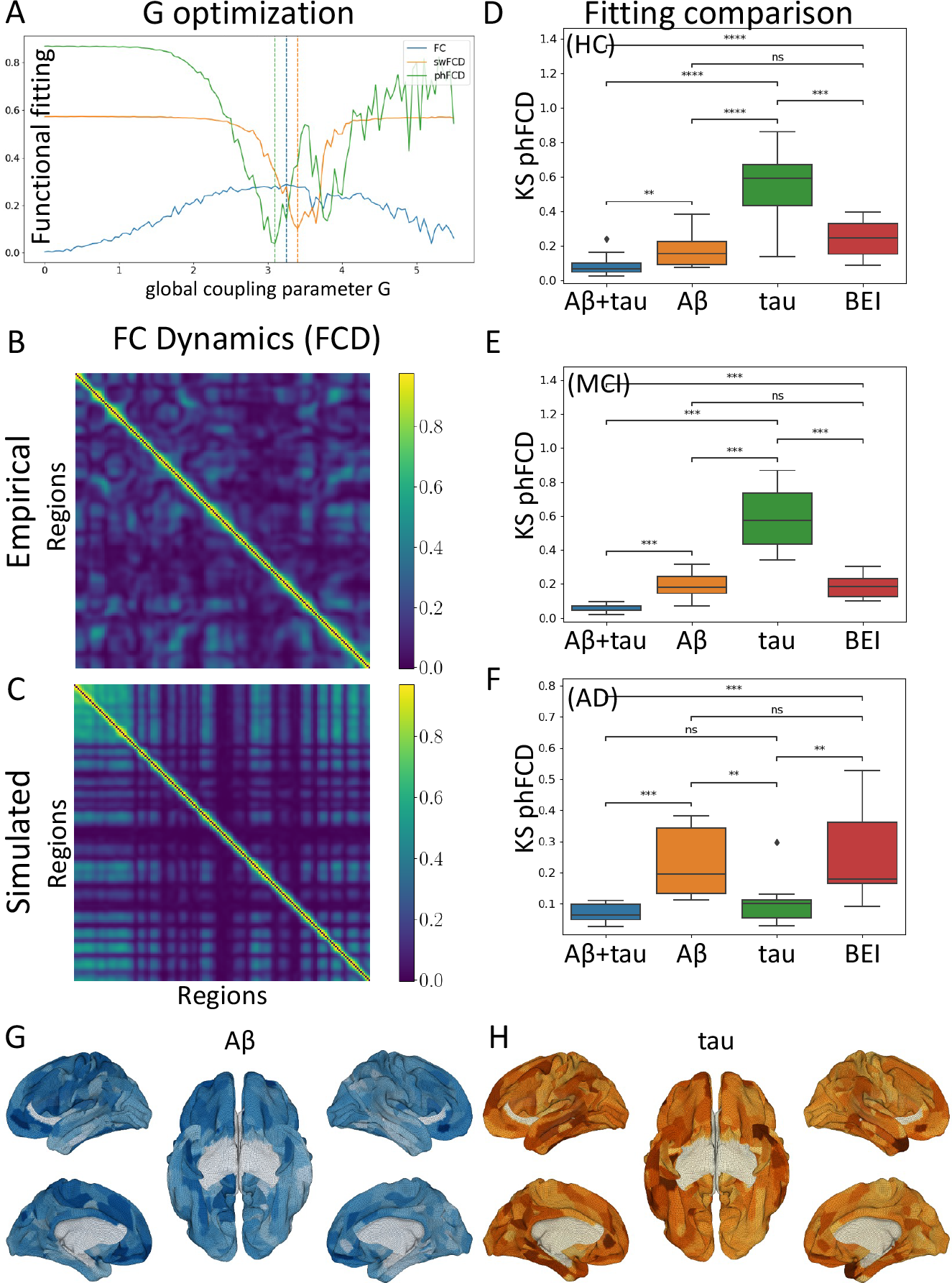
Optimization and evaluation of the model: First, using only HC subjects, the global coupling parameter *G* is found, and then the model is adjusted to minimize the distance between the empirical and simulated fMRI data, taking into account the regional burden distributions. (A) Minimization of *G* between 0 and 5.5, for Functional connectivity (FC), sliding-window Functional connectivity Dynamics (swFCD), and phase FCD (phFCD). Given their strong similarity in the results, phFCD was used for all subsequent computations. (B, C) Shows the normalized (in [0, 1]) FCD distributions for the empirical data (top) and the simulated model at the optimal result (bottom). (D, E, F): Analysis of the impact (smaller values are better) of the different burdens with respect to their impact on the phFCD (KS distance) when optimized together and in isolation, with the homogeneous state as a reference. Clearly, in all cases, the combined burden outperforms any other model. However, as can be seen, the results for AD clearly show that tau alone accounts for the vast majority of the weight of the impact on brain activity (F), while for MCI patients it is A*β* who dominates (E). For HC patients we also see a predominance of A*β*, although with less difference between the model incorporating A*β* and tau vs. A*β* in isolation (D). Example distributions of A*β* (G) and Tau burdens (H) of one subject (036_S_4430 in ADNI’s database). Colors correspond to the normalized burden of each protein.

### 2.6 Simulated BOLD signal

Once we have obtained the simulated mean field activity, we need to transform it into a BOLD signal we used the generalized hemodynamic model of Stephan et al. [37]. We compute the BOLD signal in the *k*-th brain area from the firing rate of the excitatory pools *H*^(*E*)^, such that an increase in the firing rate causes an increase in a vasodilatory signal, *s*_*k*_, that is subject to auto-regulatory feedback. Blood inflow *f*_*k*_ responds in proportion to this signal inducing changes in blood volume *v*_*k*_ and deoxyhemoglobin content *q*_*k*_. The equations relating these biophysical variables are:

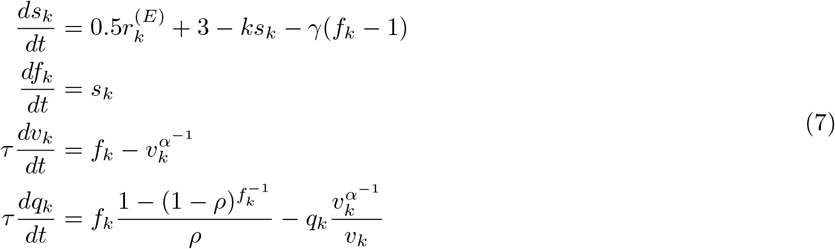

with finally

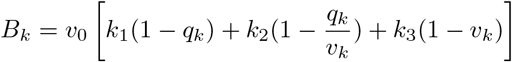

being the final measured BOLD signal.

We actually used the updated version described later on [37], which consists on introducing the change of variables 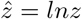, which induces the following change for *z* = *f*_*k*_, *v*_*k*_ and *q*_*k*_, with its corresponding state equation 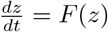, as:

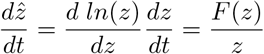

which results in 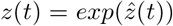 always being positive, which guarantees a proper support for these non-negative states, and thus numerical stability when evaluating the state equations during evaluation.

### 2.7 A*β*-Tau model

In our heterogeneous model, A*β* and Tau are introduced, at the formulae for the neuronal response functions, *H*^(*E,I*)^ (excitatory/inhibitory), into the gain factor 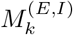 for the *k*-th area as

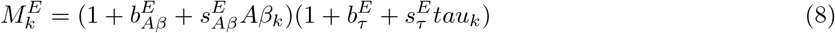

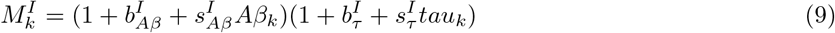

where 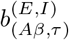 are the excitatory/inhibitory A*β* and tau bias parameters, while 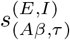 are the respective scaling factors. These are the 8 (from which actually only 6 are needed as tau only affects excitatory neurons [38], see next section) parameters that we will optimize for each subject individually.

### 2.8 Constraints

Based on previous neuroscientific experiments [4], we included constraints on the direction of effect of A*β* and tau (i.e., inhibitory vs. excitatory influence). We introduced the following constraints:

- A*β* produces inhibitory GABAergic interneuron dysfunction [6, 39], thus we can infer that 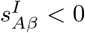.
- A*β* produces impaired glutamate reuptake [6, 39], so we can introduce the bound 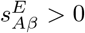.
- Tau appears to target excitatory neurons [38], so we can safely consider that 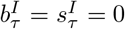.
- Tau binds to synaptogyrin-3, reducing excitatory synaptic neurotransmitter release [40], thus 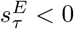.

Although the interplay between A*β* and tau is not completely known [4], but there is evidence that A*β* promotes tau by cross-seeding [41, 42], thus the cross term factors (i.e., the ones resulting from the multiplication of A*β* and tau scaling parameters) play a crucial role to elucidate the final impact of the combined burden.

### 2.9 Observables

#### edge-centric FC

The static edge-level FC is defined as the *N × N* matrix of BOLD signal correlations between brain areas computed over the entire recording period (see Figure 3). We computed the empirical FC for each human participant and for each simulated trial, as well as for the group-averages SC matrix of the healthy cohort. All empirical and simulated FC matrices were compared by computing the Pearson correlation between their upper triangular elements (given that the FC matrices are symmetric).

**Figure 3.**
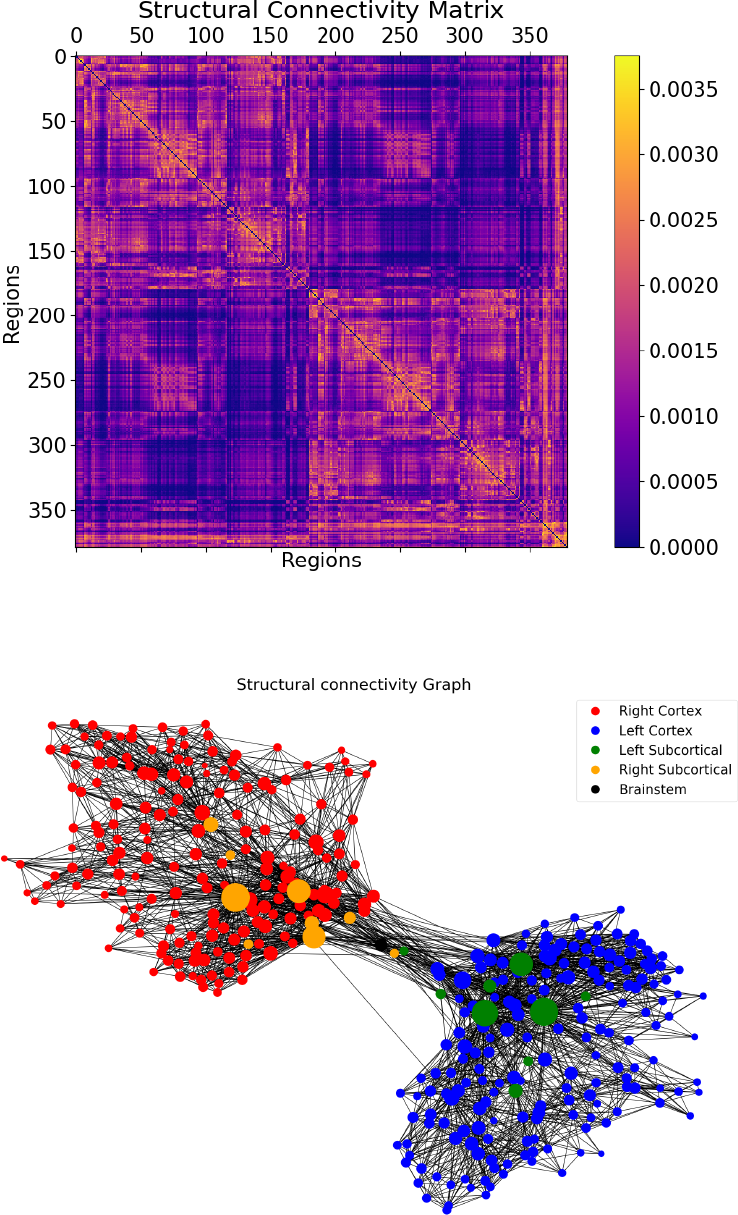
Visualization of the SC graph, in matrix form (left) and as a graph showing the strongest 5% of connections. Node positions are computed with Fruchterman and Reingold’s [52] algorithm, which assumes stronger forces between tightly connected nodes. Besides the high degree of symmetry, we can observe the laterality is kept in the graph structure (also for subcortical regions). Node size linearly represents the graph theoretical measure of structural degree for each node. As we can see, the most important hubs are in the subcortical regions.

#### swFCD

The most common and straightforward approach to investigate the temporal evolution of FC is the sliding-window FC dynamics (swFCD) [43]. This is achieved by calculating the correlation matrix, *FC*(*t*), restricted to a given time-window (*t − x* : *t* + *x*), and successively shifting this window in time resulting in a time-varying *FC*_*NxNxT*_ matrix (where *N* is the number of brain areas and *T* the number of time windows considered). Here, we computed the FC over a sliding window of 30 TRs (corresponding approximately to 1.5 minutes) with incremental shifts of 3 TRs. This FCD matrix is defined so that each entry, (*FCD*(*t*_*x*_, *t*_*y*_)) corresponds to the correlation between the FC centered at times *t*_*x*_ and the FC centered at *t*_*y*_. In order to compare quantitatively the spatio-temporal dynamical characteristics between empirical data and model simulations, we generate the distributions of the upper triangular elements of the FCD matrices over all participants as well as of the FCD matrices obtained from all simulated trials for a given parameter setting. The similarity between the empirical and model FCD distributions is then compared using the KS distance, *D*_*KS*_, allowing for a meaningful evaluation of model performance in predicting the changes observed in dynamic resting-state FC. However, the fundamental nature of the swFCD technique implies the choice of a fixed window length, which limits the analysis to the frequency range below the window period, so the ideal window length to use remains under debate [44].

#### phFCD

In an attempt to overcome the limitations of the sliding-window analysis, a few methods were proposed to estimate the *FC*(*t*) at the instantaneous level. For instance, phase Functional Connectivity Dynamics (*phFCD*) consists in computing the phase coherence between time series at each recording frame [21]. In brief, the instantaneous BOLD phase of area *n* at time *t, θ*_*n*_(*t*), is estimated using the Hilbert transform. Given the phase, the angle between two BOLD signals is given by their absolute phase difference: Θ_*np*_ = |*θ*_*n*_(*t*) *− θ*_*p*_(*t*)|. Then, the *phFCD*(*t*) between a pair of brain areas *n* and *p* is calculated as:

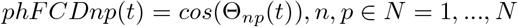

with *N* the number of brain regions considered in the parcellation used. To compare two phFCD matrices among themselves, e.g., a simulated and an empirical one, again the KS distance is usually used.

### 2.10 Full Optimization

To efficiently optimize the 6-dimensional function described before for the three bias and scaling values, a simple local optimization-based approach such as conjugate gradients cannot be used, as this is a (usually) ill-posed problem with a global minimum surrounded by many local minima. Instead, we need to resort to a global optimization algorithm. In our case, mwe used a Bayesian minimization algorithm using Gaussian Processes (GP), which approximates the function using a multivariate Gaussian. In particular, our implementation uses the *gp minimize* method from the *scikit-optimize* Python library^[5]^. At its core, the method approximates the objective function with a Gaussian process, assuming that the values follow a multivariate Gaussian. The covariance of the function values is given by a GP kernel between the parameters. With this information, the algorithm chooses the next parameter to evaluate by selecting the acquisition function over the Gaussian prior. The error measure used was the KS distance between the empirical BOLD signal and the average over a number of trials (10 in our case) of the simulated signal, and we let the function run for 100 iterations. In all cases, the results moved significantly away from the priors.

## 3 Results

We used diffusion MRI to generate the Structural Connectomes of 17 healthy control (HC) subjects, 9 mild cognitive impairment (MCI) subjects, and 10 subjects with Alzheimer’s Disease (AD) from ADNI, which are mostly the same participants as those used by Stefanovski et al. [15] and Triebkorn et al. [16]. See Table 2 and Figure 3.

**Table 2.** Epidemiological information of the population used in this study.

| Diagnosis | n (female) | Mean age | $\sigma$ | Min. age | Max. age | Mean MMSE | $\sigma_{MMSE}$ | Min. MMSE | Max. MMSE |
| --- | --- | --- | --- | --- | --- | --- | --- | --- | --- |
| AD | 10 (5) | 72.0 | 9.6 | 55.9 | 86.1 | 21.3 | 6.8 | 9 | 30 |
| HC | 17 (10) | 70.8 | 4.3 | 63.1 | 78.0 | 29.3 | 0.7 | 28 | 30 |
| MCI | 9 (3) | 68.8 | 5.8 | 57.8 | 76.6 | 27.4 | 1.5 | 25 | 30 |

Given these cohorts, we used the G∗Power [45] software to conduct statistical power calculations based on a two-group Wilcoxon-Mann-Whitney test, with significance level *α* = 0.05 and power 1 *− β* = 0.8. Assuming a standard deviation *σ* = 0.05 (a reasonable assumption given our results below), we obtained that the minimum effect size we would be able to discern in this setting would be *d* = 1.1, which implies that the minimum detectable difference between the means of the control population and any of the other two would be around 0.055.

### 3.1 Fitting the Homogeneous Model

As a first step, we evaluated the capability of the homogeneous BEI model to reproduce the empirical properties of resting-state FC data. To this end, we fitted the global coupling parameter, *G*, without considering heterogeneity by setting all regional gain parameters *M*_(*E,I*)_ = 1 [18]. Then, we evaluated the ability of the model to reproduce three different properties of empirical resting-state fMRI recordings: edge-level static FC, swFCD, and phFCD (see Methods for further details.) The results of this analysis are shown in Figure 2A. To remove differences across subjects related to age, we considered averaged values across subjects over the healthy control group and took an equivalent number of simulated trials with the same duration as the human experiments (see Methods). Following previous research [19] fitting the phFCD better captures the spatiotemporal structure of the fMRI data, being a stronger constraint on the model. Indeed, where FC fits are consistently high across a broad range of *G* values, phFCD yields a clear global optimum at *G* = 3.1. Thus, we choose to use phFCD for all further analysis.

### 3.2 Introducing A*β* and tau heterogeneity

Once the global coupling parameter has been found, we can introduce the regional heterogeneity in the distributions of A*β* and tau, and study how their introduction leads to a better representation of neural dynamics, i.e., improves the fitting of phFCD. Spatial maps for each form of protein burden used in our modeling are shown in Figures 2G (for A*β*) and 2H (for tau) for one particular individual. For some individuals, (mainly HC subjects, e.g., as subject 003 S 6067 in the ADNI database, with *ρ* = 0.92, *p <* 0.001) the A*β* and tau distributions are strongly correlated, while for others the two maps show a weaker correlation (e.g., subject 036_S_4430, with *ρ* = 0.10, *p* = 0.04.) This observation indicates that each protein burden introduces a different form of biological heterogeneity to the benchmark BEI model, and thus should be modeled separately in our simulations.

We introduce these kinds of heterogeneity by modulating the regional gain functions *M*_(*E,I*)_ at the optimal working point of the homogeneous BEI model found at the previous stage (*G* = 3.1), through the bias and scaling parameters introduced above, denoted 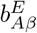 and 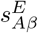 for A*β*, and 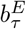 and 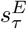 for tau, all for the excitatory case, and similarly for the inhibitory case with superscript *I*. We perform a search in parameter space with constraints introduced from experimental observations, see Section 2.8, to find the optimal working point for the two protein burdens simultaneously, which results in an 8-degree of freedom optimization, which is reduced to six degrees due to the constraints. For the optimization, we used **Bayesian optimization algorithm using Gaussian Processes**, see Section 2.10. We can also perform a simplified search, limited to the two-variable 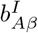 and 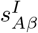 space, i.e., the inhibitory bias and scaling of the A*β* influence on inhibitory neuron parameters (Equation 9). In this case, the 2D optimization results show a decreasing the neuronal activity with increasing A*β* concentration, confirming previous results [15]. On average, for each group of subjects, we got the results shown numerically in Table 3.

**Table 3.** Resulting averaged parameters from the optimization procedure. In parenthesis, the respective standard deviations.

| Cohort | $b_{A\beta}^E$ | $s_{A\beta}^E$ | $b_\tau^E$ | $s_\tau^E$ | $b_{A\beta}^I$ | $s_{A\beta}^I$ |
| --- | --- | --- | --- | --- | --- | --- |
| AD | 0.2 (0.5) | 2.3 (1.2) | -0.4 (0.6) | -2.6 (0.8) | 0.2 (0.6) | -2.5 (0.8) |
| MCI | 0.4 (0.7) | 1.7 (1.5) | -0.5 (0.5) | -2.8 (0.7) | -0.1 (0.8) | -2.1 (1.2) |
| HC | 0.1 (0.8) | 1.7 (0.9) | -0.5 (0.6) | -2.8 (1.0) | 0.3 (0.6) | -3.1 (1.0) |

These results can be seen visually in Figure 4. This figure shows that there is a clear regime in which all three empirical properties are fitted well by the model, particularly for the values shown above, where a fitting of phFCD of 0.13 is achieved for the AD subjects, while the reference homogeneous value is equal to 0.5.

**Figure 4.**
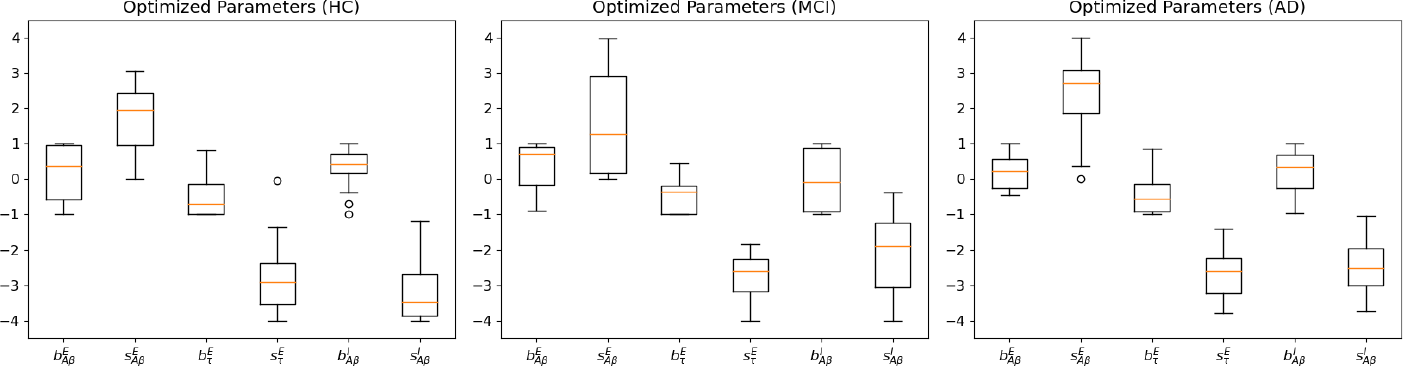
Parameter values found after the optimization stage for HC, MCI and AD subjects. Observe that all 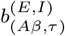, the excitatory/inhibitory A*β* and tau bias parameters, have negligible values, while the scaling parameters 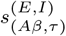 present non-null values. Of note, the p-values between the different scaling parameters across the cohorts are different in a moderately significant way (*p <* 0.03), remarkably between HC and AD, but usually not between MCI and AD. In these plots, boxes extend from the lower to upper quartile values of the data, adding an orange line at the median. Also, whiskers are used to show the range of the data, extending from the box.

### 3.3 Analysis of burden impact

For the optimal parameter values resulting from model fitting, we simulated each dynamical model 10 times for each subject to account for the inherently stochastic nature of the models and compute the respective measures of model fit. Figure 5 shows the distributions of fit statistics across runs for the homogeneous and the heterogeneous model for the different cohorts. In addition, we show results for a null ensemble of models in which the regional burden values were spatially shuffled to generate surrogates with the same spatial autocorrelation as the empirical data. Across the benchmark property to which the data were fitted –—phFCD-––, the models taking into account the regional burden heterogeneity perform better than the homogeneous model (all pass the Mann-Whitney U rank test on two *independent samples* with *p <* .0005). We also find a consistent gradient of performance across all benchmarks, with the heterogeneous model performing best, and the homogeneous model showing the poorest performance. For each benchmark metric, the performance of the heterogeneous model was better than all other models (in all cases *p <* .06). Also, it must be noted that the differences in fit statistics between models are significant, as shown in Figure 5. For example, for the AD cohort, the correlation of the median phase FCD between the fitted model and empirical data showed *r <* 0.1 for the heterogeneous model, and *r* ≈ 0.2 for the BEI model. In all subject groups, the difference between these two models is clear, with *p* < 0.0005. In all reported results we used a Mann-Whitney-Wilcoxon test two-sided with Benjamini-Hochberg correction (p-value annotation legend: ns: *p* <= 1.00*e* + 00, *: 1.00*e −* 02 *< p* <= 5.00*e −* 02, ^**^: 1.00*e −* 03 < *p <*= 1.00*e −* 02, ^***^: 1.00*e −* 04 < *p <*= 1.00*e −* 03, ^****^: *p* <= 1.00*e −* 04).

**Figure 5.**
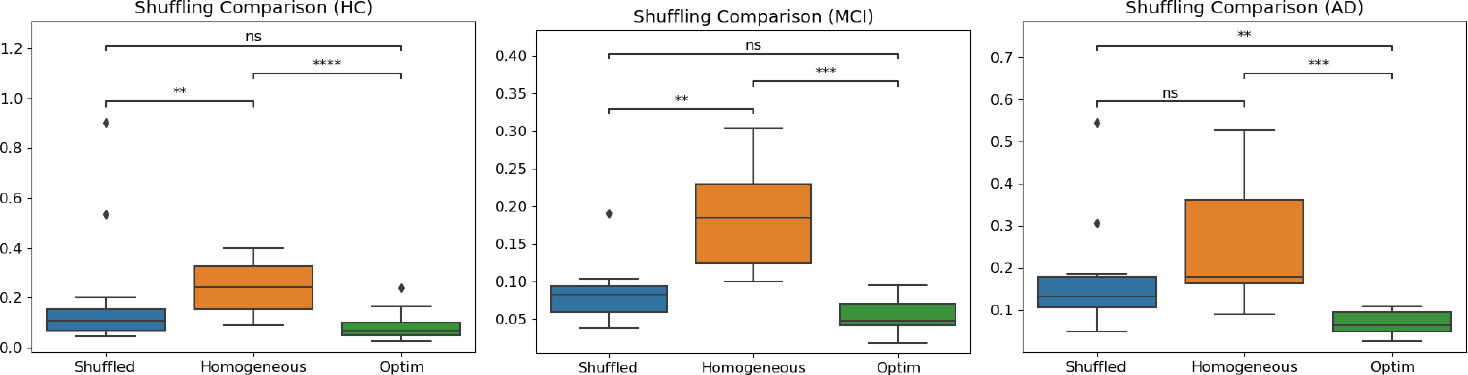
Comparison between the homogeneous model, the optimum result obtained with the heterogeneous model (optim), and the same parameter values but with shuffled burdens. As can be seen, the differences in fit statistics between models are significant. In particular, for the AD cohort, the median phFCD correlation between model and data showed *r <* 0.1 for the these two models is clear, with *p <* 0.0005. heterogeneous model, and *r ≈* 0.2 for the BEI model. In all subject groups, the difference between these two models is clear, with *p* < 0.0005.

Finally, we performed an analysis comparing the impact of each type of burden, in isolation or together, onto the simulation results. In Figures 2D-2F we can see these results for the different cohorts, for A*β* and tau, A*β* alone, tau alone and finally the homogeneous BEI model, added for reference. As we can see, with respect to the homogeneous model, the best performance is systematically obtained by the combined action of both A*β* and tau, giving a value with *p* < 0.0004 in all cases. However, for each cohort, each protein shows to play a different role in the development of the disease. For AD subjects, the effect of A*β* on the optimal combined result is small, with a *p* < 0.0005, while the influence of tau alone has a *p* value that does not allow us to distinguish between its effect and the combined effect of both proteins (*p* = 0.172), implying a clear dominance of tau over A*β* in this stage of the disease. Also, with respect to the homogeneous BEI model, tau presents *p* < 0.005, while A*β* alone shows a much higher value (*p* = 0.339), not allowing us to clearly distinguish between these two models. In the case of the MCI cohort, in Figure 2E, we can observe that the effect of A*β* alone clearly gives the major contribution to the final combined fitting, rather than tau, with a *p* < 0.0003 between all cases. Finally, in the HC case in Figure 2D, the effects of the A*β* and tau proteins are close to the homogeneous BEI model, with A*β* presenting a somewhat higher prevalence than tau. However, it is noticeable that the differences between this case and the previous one are small, showing that A*β* already plays an important role even in HC subjects.

### 3.4 2D A*β* Optimization

We can use our model to verify the results by Stefanovski et al. [15] by limiting our analysis to the parameters of A*β* at the inhibitory level (i.e., the inhibitory bias 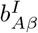 and scaling 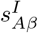 parameters only, defined in Equation 9). This way, we can replicate, up to a certain degree, the results from that paper, being limited by the fact that we use a different model, based on the BEI model instead of the Jansen-Rit model [46]; a different expression for the burden, i.e., a linear approximation instead of a sigmoid; different units, etc. See Figure 4. By analyzing the obtained data at the optimal fit, the same behavior of decreasing the neuronal activity of inhibitory neurons with the scaling parameter 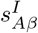, corresponding to an increase in A*β* concentration, can be observed, as shown in Figure 6.

**Figure 6.**
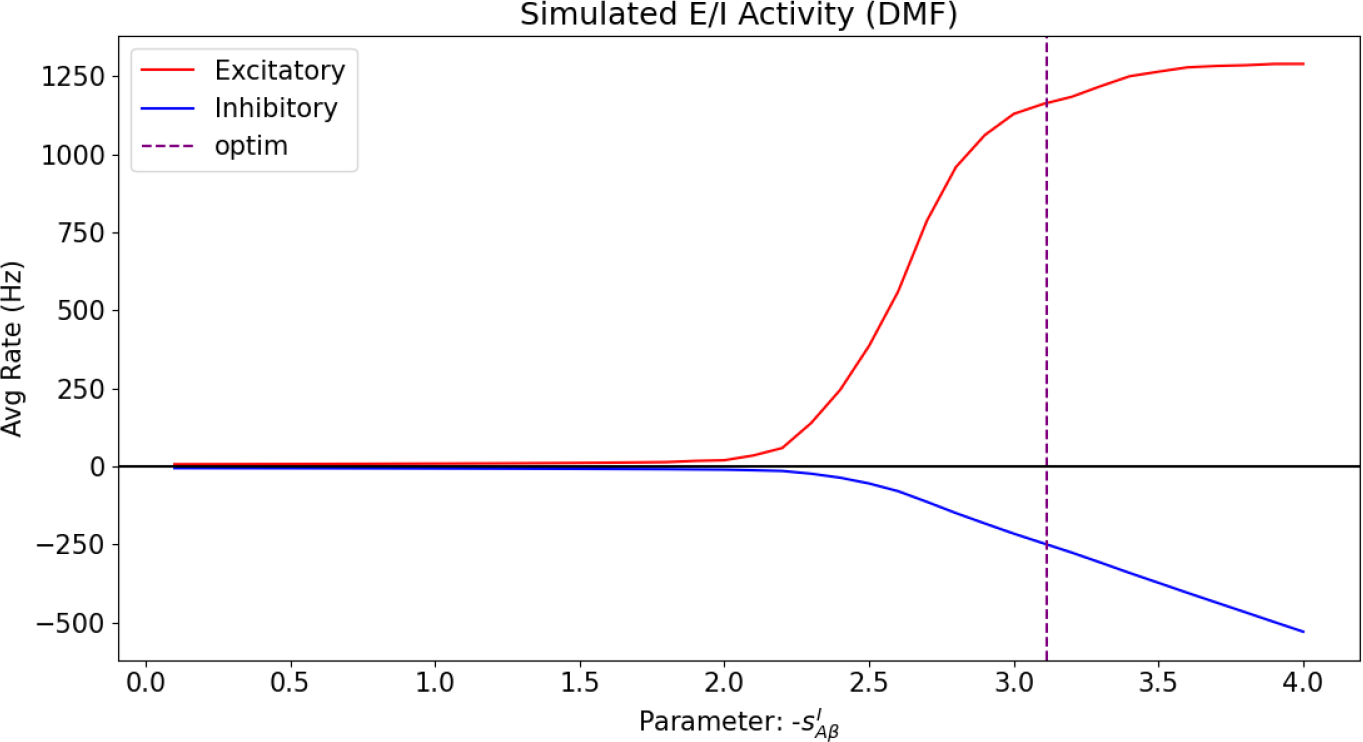
Excitatory and inhibitory mean firing rates as a function of the A*β* inhibitory scaling 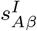, with all the other parameters of the model at the (averaged) fitted optimum values. For the purpose of clarity, the horizontal axis for the scaling has been taken as absolute values, to illustrate the behavior with increasing A*β* loads. The vertical axis shows the firing rates of both excitatory and inhibitory populations. It can be clearly seen that the net effect of the burden is to increase the overall region firing rate, measured at the excitatory population. For the sake of clarity, the inhibitory firing rate has been vertically inverted (negated) to show their decreased effect on the excitatory population, thus confirming previous findings [15]. The vertical discontinuous line shows the optimum found for 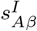 .

## 4 Discussion

In this paper we studied the influence of the regional variability of two pathological proteins, namely A*β* and tau, on cortical activity and E/I balance in the context of AD using whole-brain dynamic modeling, which allowed us to disentangle separate and synergistic effects of these two proteins in-silico. The incorporation of such heterogeneous patterns of neuropathology into whole-brain models of neuronal dynamics has been made possible by the availability of in-vivo quantitative PET imaging. We have shown that the heterogeneous model, which incorporates regional information on both types of neuropathological burdens more faithfully reproduces empirical properties of dynamic brain activity than the model with fixed and homogeneous parameters. Our findings highlight the central role of both types of burden in disturbing the E/I balance, supporting the hypothesis of hyperexcitation in AD. Regarding the individual influence of A*β* and tau on brain activity, our results have shown a dominance of A*β* influence on neural dynamics in earlier stages of AD (i.e., MCI) and even in healthy controls, while tau plays a larger role in later stages. These key findings highlight the prominent role of these pathological proteins in contributing to the abnormal brain activity patterns in the course of AD [47].

### 4.1 How does burden heterogeneity shape neuronal dynamics?

We introduced burden heterogeneity into our dynamical model by modifying the regional excitability of neural population activity. We achieved this by modifying each brain region’s gain response function *M*_*i*_ of inhibitory and excitatory populations, i.e., the net excitability of the according population. This was done, in accordance with previous works exploring the effect of regional parameters on E/I balance [19], thus focusing on how the interaction of neuronal populations contributes to neuronal dynamics (i.e., FC or FCD) and their relative impacts over time. Our approach is different from the work by Stefanovski et al. [15], where the A*β* burden was used to modulate regional E/I balance by negatively modulating the inhibitory time constant, slowing down synaptic transmission and thus increasing excitatory activity and producing a higher output of the pyramidal cell populations, resulting in a local hyperexcitation with high A*β* loads. However, our approach, when limited to the effect of A*β* in the early stages of the disease, results in the same behavior of neural populations as a function of A*β*, similarly resulting in a net increase of the excitatory activity with increased A*β* burden. There are other approaches available to introduce heterogeneity, such as an adjustment of the inter-node connectivity to fit empirical and simulated FCs [48]; or variations of within- and inter-area connectivity [49]. However, based on the empirical evidence that the interplay of both burdens, A*β* and tau, severely disrupt normal neuronal function, we decided to model their direct effect on the E/I balance.

In this paper we have chosen to incorporate heterogeneity into the model by modulating population gain response functions *H*^(*E,I*)^. Here, adjusting the gain function parameter *M*_*i*_ allows us to demonstrate how local variations in the E/I balance will affect the net excitability of the population. We thus assume that changes in regional gain are the common final pathways of different neuropathology-related pathomechanisms which might have an influence on neuronal populations, i.e., result in realistic representation of direct effects of A*β* and tau and also associated processes (i.e., non-direct effects).

In particular, we introduced regional variations of *M*_*i*_ as the product of linear terms consisting of a constant (bias), and a scaling factor. This introduced eight degrees of freedom, which we could narrow down to six degrees of freedom by introducing constraints to the direction of effect based on previous research [4]. In sum, our model was created based on assumptions that A*β* leads to GABAergic interneuron dysfunction and impaired glutamate reuptake, while tau leads to reduced synaptic neurotransmitter release in excitatory cells. This hypothesis-driven amount of degrees is substantially less than used in other models [49, 48], making a fast parameter optimization feasible, while ensuring sufficient biological realism. Furthermore, in all cases, the bias parameters for the different burdens (Figure 4) were approximately 0, thus indicating that the influence of the bias parameters with respect to the homogeneous model can be ignored, further reducing computational complexity. The respective scaling parameters take non-negligible values, showing a linear relationship between A*β* and tau on neural dynamics. We used Bayesian optimization using Gaussian Processes (see Methods) to address the challenge of multiple local minima that could trap traditional optimization methods.

### 4.2 Evaluating A*β* and tau impact

A large body of scientific literature focused on linking global and local brain dynamics to individual differences in cognitive performance scores [12] and showed that patients with AD and MCI show less variation in neuronal connectivity during resting-state, and even presented benchmarks for predictive models based on resting-state fMRI, revealing biomarkers of individual psychological or clinical traits [13]. However, the pattern of neuronal connectivity alterations has been incompletely understood. More recent work focused on the effect of A*β* on hyperexcitability, and how A*β* modulates regional E/I balance, resulting in local hyperexcitation in brain regions with high loads of A*β* [15]. To our knowledge, no prior study has evaluated both types of neuropathological burden, A*β* and tau, simultaneously, linking neuropathological data with dynamic whole-brain modeling.

As explained in the Methods Section, we compared the impact of each type of burden, in isolation or interaction, onto neural dynamics. We found that the model fitting optimum is systematically obtained by the interaction of both burdens, underlining the interaction of both proteins in disturbing neural activity. Also, we have found that for each condition (i.e., HC, MCI or AD), each protein has a different impact on the brain dynamics. In the case of AD, A*β* has a small impact on the combined result, while tau alone had almost all of the impact, showing its dominance over A*β* in regard to generating abnormal brain dynamics. Also, in comparison to the homogeneous BEI model, we observed that tau is clearly distinguishable, but A*β* is not. Taken together, these results imply that we cannot distinguish between the effect on the brain activity of both proteins together vs. the effect of tau alone, while the effect of A*β* is clearly distinguishable from the combined effect. As a consequence, this allows us to conclude that the impact of tau in the late stage of the disease (AD) is clearly dominant over A*β*. In contrast, in MCI, the influence of A*β* alone is clearly dominant over tau, see Figure 2E. Finally, when studying the effect of both proteins in HC, we can observe that the effect of the A*β* and tau proteins is close to the homogeneous BEI model, with A*β* presenting a relatively higher influence than tau. The influence of A*β* both in MCI patients as well as in HC shows that A*β* leads to a measurable change in brain dynamics in elderly people, independent of existing cognitive impairment. However, we acknowledge that on a pathophysiological level, there is a strong interplay between A*β* and tau, and further (causal) research is needed to clearly discern the role each protein plays in the generation of neuronal dysfunction. Despite our findings from model fitting, we acknowledge that we only observe the current influence of A*β* vs. tau in different disease stages in a cross-sectional cohort. Longitudinal examinations might also replicate the abundant evidence in the literature [4] that both proteins interplay a toxic feedback loop, which is ultimately responsible (perhaps among other factors) for the development of the disease.

Our analysis based on furthermore shows that edge-level measures of static FC offer loose constraints for model optimization, showing comparably high fit statistics across a broad range of values of the global coupling parameter. In contrast, fitting to dynamical functional connectivity shows a clear optimum, mirroring similar results reported previously [19]. We can conclude that fitting models to both static and dynamic properties is thus important for identifying an appropriate working point for each model.

Across all these properties, we observe that the model that incorporates the heterogeneous burden loads provides a better match to the data than the homogeneous BEI model, which does not incorporate a fitting of the gain response function of inhibitory and excitatory populations to the data. This shows that constraining regional heterogeneity by the protein burdens yields a more faithful replication of brain dynamics, as measured by empirical phFCD. The superiority of our model using heterogeneous, empirically estimated parameters, suggests that regional heterogeneity plays a significant role in shaping the effects of Alzheimer’s disease on spontaneous BOLD-dynamics. As we already mentioned, it must be noted that the differences in fit statistics between models are significant. These results suggest that these empirical fit statistics have good capacity to tease apart dynamical differences between models, which gives the opportunity to disentangle the influence of different pathomechanisms in vivo.

## 5 Limitations

In our implementation, we used SC matrices derived from DWI. However, as many factors such as myelination and diameter impact the conductivity of white matter tracts, it may be a confounding factor that coupling between different brain areas is not affected by this.

It is important to mention that, in our study, we included sub-cortical regions, which are particularly susceptible to off-target binding of the AV-1451 tracer, which may introduce a potential confound. For this study we did not have controlled the images for off-target binding. Recent studies show that, besides Tau tangles, AV-1451 also binds on neuromelanin, melanin and other blood products [50]. This is a genuine restriction of the imaging method (as many PET tracers have one) and we are not aware of a standard that corrects for this phenomenon. Moreover, it is kind of controversially discussed whether this is always off-target binding or detecting tangles that other methods are not aware of [51]. In any case, it can be argued that the typical finding of this off-target bidding is affecting AD, MCI and controls to the same amount, which implies this is something that cannot cause artificial group differences and can therefore be safely ignored in the context of this study.

## 6 Conclusion

In summary, in this paper, we have presented a whole-brain dynamic model connecting the main protein burdens, namely A*β* and tau, in different stages of AD and in HC. Our results not only reproduce previous research regarding E/I imbalance in AD, and also shed further light on the relative impact of each type of burden during different disease stages, opening new avenues to focus research efforts. As a general conclusion, our study shows that theory-driven whole-brain modeling enables us to do research on disease mechanisms in-silico and to empirically compare competing hypotheses against each other and thus complements data-driven modeling such as machine learning. Thus, whole-brain modeling can incorporate sufficient biological realism to contribute to improved diagnostic procedures (i.e., enable the use of fMRI for diagnosis) to discover new therapies (e.g., by simulating novel treatments).

## Declarations

### Ethics approval and consent to participate

Ethical approval was obtained by ADNI sites and written informed consent was collected from all participants. No further consent was necessary.

### Availability of data and materials

All code for implementing computational models and reproducing our results will be available at the first author’s repository: https://github.com/dagush/WholeBrain Data used in preparation of this article were obtained from the Alzheimer’s Disease Neuroimaging Initiative (ADNI) database (`adni.loni.usc. edu`). As such, the investigators within the ADNI contributed to the design and implementation of ADNI and/or provided data but did not participate in analysis or writing of this report. A complete listing of ADNI investigators can be found at: https://adni.loni.usc.edu/wp-content/uploads/how_to_apply/ADNI_Acknowledgement_List.pdf.

### Authors’ contributions

**Gustavo Patow**: Conceptualization, Formal analysis, Software, Writing – original draft, Writing – review & editing. **Leon Stefanovski**: Data Curation, Writing – review & editing. **Petra Ritter**: Data Curation, Writing – review & editing. **Gustavo Deco**: Conceptualization, Writing – review & editing. **Xenia Kobeleva**: Conceptualization, Writing – review & editing.

## Acknowledgements

Data collection and sharing for this project was funded by the Alzheimer’s Disease Neuroimaging Initiative (ADNI) (National Institutes of Health Grant U01 AG024904) and DOD ADNI (Department of Defense award number W81XWH-12-2-0012). ADNI is funded by the National Institute on Aging, the National Institute of Biomedical Imaging and Bioengineering, and through generous contributions from the following: AbbVie, Alzheimer’s Association; Alzheimer’s Drug Discovery Foundation; Araclon Biotech; BioClinica, Inc.; Biogen; Bristol-Myers Squibb Company; CereSpir, Inc.; Cogstate; Eisai Inc.; Elan Pharmaceuticals, Inc.; Eli Lilly and Company; EuroImmun; F. Hoffmann-La Roche Ltd and its affiliated company Genentech, Inc.; Fujirebio; GE Healthcare; IXICO Ltd.; Janssen Alzheimer Immunotherapy Research & Development, LLC.; Johnson & Johnson Pharmaceutical Research & Development LLC.; Lumosity; Lundbeck; Merck & Co., Inc.; Meso Scale Diagnostics, LLC.; NeuroRx Research; Neurotrack Technologies; Novartis Pharmaceuticals Corporation; Pfizer Inc.; Piramal Imaging; Servier; Takeda Pharmaceutical Company; and Transition Therapeutics. The Canadian Institutes of Health Research is providing funds to support ADNI clinical sites in Canada. Private sector contributions are facilitated by the Foundation for the National Institutes of Health (www.fnih.org). The grantee organization is the Northern California Institute for Research and Education, and the study is coordinated by the Alzheimer’s Therapeutic Research Institute at the University of Southern California. ADNI data are disseminated by the Laboratory for Neuro Imaging at the University of Southern California.

The authors would like to thank Javier Palarea Albaladejo for his invaluable help with the statistical analysis.

## Abbreviations

(SC): structural connectome
(A*β*): amyloid-beta
(AD): Alzheimer’s Disease
(HC): healthy controls
(MCI): mild cognitive impairment
(BEI): Balanced Excitation-Inhibition
(FC): functional connectivity
(swFCD): sliding-window functional connectivity dynamics
(phFCD): phase functional connectivity dynamics
(KS): Kolmogorov-Smirnov distance
(ADNI): Alzheimer’s Disease Neuroimaging Initiative

## Footnotes

[1] https://surfer.nmr.mgh.harvard.edu/fswiki/FreeSurferMethodsCitation

[2] https://www.humanconnectome.org/software/connectome-workbench

[3] http://www.mrtrix.org

[4] https://mrtrix.readthedocs.io/en/latest/dwi_preprocessing/dwipreproc.html

[5] https://scikit-optimize.github.io/stable/modules/generated/skopt.gp_minimize.html

## References

1. Stefanovski, L., Meier, J.M., Pai, R.K., Triebkorn, P., Lett, T., Martin, L., Bülau, K., Hofmann-Apitius, M., Solodkin, A., McIntosh, A.R., Ritter, P.: Bridging scales in alzheimer’s disease: Biological framework for brain simulation with the virtual brain. Frontiers in Neuroinformatics 15 (2021). doi:10.3389/fninf.2021.630172

2. Jack, C.R., Bennett, D.A., Blennow, K., Carrillo, M.C., Dunn, B., Haeberlein, S.B., Holtzman, D.M., Jagust, W., Jessen, F., Karlawish, J., Liu, E., Molinuevo, J.L., Montine, T., Phelps, C., Rankin, K.P., Rowe, C.C., Scheltens, P., Siemers, E., Snyder, H.M., Sperling, R., Elliott, C., Masliah, E., Ryan, L., Silverberg, N.: NIA-AA research framework: Toward a biological definition of alzheimer’s disease. Alzheimer’s & Dementia 14(4), 535–562 (2018). doi:10.1016/j.jalz.2018.02.018

3. Alexander, G.C., Karlawish, J.: The problem of aducanumab for the treatment of alzheimer disease. Annals of Internal Medicine 174(9), 1303–1304 (2021). doi:10.7326/m21-2603

4. Busche, M.A., Hyman, B.T.: Synergy between amyloid-β and tau in alzheimer’s disease. Nature Neuroscience 23(10), 1183–1193 (2020). doi:10.1038/s41593-020-0687-6

5. Lock, M.: The Alzheimer Conundrum : Entanglements of Dementia and Aging. Princeton University Press, Princeton (2013)

6. Palop, J.J., Mucke, L.: Network abnormalities and interneuron dysfunction in alzheimer disease. Nature Reviews Neuroscience 17(12), 777–792 (2016). doi:10.1038/nrn.2016.141

7. Maestú, F., de Haan, W., Busche, M.A., DeFelipe, J.: Neuronal excitation/inhibition imbalance: core element of a translational perspective on alzheimer pathophysiology. Ageing Research Reviews 69, 101372 (2021). doi:10.1016/j.arr.2021.101372

8. Chang, C.-W., Evans, M.D., Yu, X., Yu, G.-Q., Mucke, L.: Tau reduction affects excitatory and inhibitory neurons differently, reduces excitation/inhibition ratios, and counteracts network hypersynchrony. Cell Reports 37(3), 109855 (2021). doi:10.1016/j.celrep.2021.109855

9. Bi, D., Wen, L., Wu, Z., Shen, Y.: GABAergic dysfunction in excitatory and inhibitory (e/i) imbalance drives the pathogenesis of alzheimer’s disease. Alzheimer’s & Dementia 16(9), 1312–1329 (2020). doi:10.1002/alz.12088

10. Petrache, A.L., Rajulawalla, A., Shi, A., Wetzel, A., Saito, T., Saido, T.C., Harvey, K., Ali, A.B.: Aberrant excitatory–inhibitory synaptic mechanisms in entorhinal cortex microcircuits during the pathogenesis of alzheimer’s disease. Cerebral Cortex 29(4), 1834–1850 (2019). doi:10.1093/cercor/bhz016

11. Lauterborn, J.C., Scaduto, P., Cox, C.D., Schulmann, A., Lynch, G., Gall, C.M., Keene, C.D., Limon, A.: Increased excitatory to inhibitory synaptic ratio in parietal cortex samples from individuals with alzheimer’s disease. Nature Communications 12(1) (2021). doi:10.1038/s41467-021-22742-8

12. Zimmermann, J., Perry, A., Breakspear, M., Schirner, M., Sachdev, P., Wen, W., Kochan, N.A., Mapstone, M., Ritter, P., McIntosh, A.R., Solodkin, A.: Differentiation of alzheimer’s disease based on local and global parameters in personalized virtual brain models. NeuroImage: Clinical 19, 240–251 (2018). doi:10.1016/j.nicl.2018.04.017

13. Dadi, K., Rahim, M., Abraham, A., Chyzhyk, D., Milham, M., Thirion, B., Varoquaux, G.: Benchmarking functional connectome-based predictive models for resting-state fmri. NeuroImage 192, 115–134 (2019). doi:10.1016/j.neuroimage.2019.02.062

14. Demirtaş, M., Burt, J.B., Helmer, M., Ji, J.L., Adkinson, B.D., Glasser, M.F., Essen, D.C.V., Sotiropoulos, S.N., Anticevic, A., Murray, J.D.: Hierarchical heterogeneity across human cortex shapes large-scale neural dynamics. Neuron 101(6), 1181–119413 (2019). doi:10.1016/j.neuron.2019.01.017

15. Stefanovski, L., Triebkorn, P., Spiegler, A., Diaz-Cortes, M.-A., Solodkin, A., Jirsa, V., McIntosh, A.R., Ritter, P.,, f.t.A.D.N.I.: Linking molecular pathways and large-scale computational modeling to assess candidate disease mechanisms and pharmacodynamics in alzheimer’s disease. Frontiers in Computational Neuroscience 13, 54 (2019). doi:10.3389/fncom.2019.00054

16. Triebkorn, P., Stefanovski, L., Dhindsa, K., Diaz-Cortes, M.-A., Bey, P., Bülau, K., Pai, R., Spiegler, A., Solodkin, A., Jirsa, V., McIntosh, A.R., and, P.R.: Brain simulation augments machine-learning–based classification of dementia. Alzheimer’s & Dementia: Translational Research & Clinical Interventions 8(1) (2022). doi:10.1002/trc2.12303

17. Deco, G., Jirsa, V.K.: Ongoing cortical activity at rest: Criticality, multistability, and ghost attractors. Journal of Neuroscience 32(10), 3366–3375 (2012). doi:10.1523/jneurosci.2523-11.2012

18. Deco, G., Ponce-Alvarez, A., Hagmann, P., Romani, G.L., Mantini, D., Corbetta, M.: How local excitation–inhibition ratio impacts the whole brain dynamics. Journal of Neuroscience 34(23), 7886–7898 (2014). doi:10.1523/JNEUROSCI.5068-13.2014. http://www.jneurosci.org/content/34/23/7886.full.pdf

19. Deco, G., Kringelbach, M.L., Arnatkeviciute, A., Oldham, S., Sabaroedin, K., Rogasch, N.C., Aquino, K.M., Fornito, A.: Dynamical consequences of regional heterogeneity in the brain’s transcriptional landscape. Science Advances 7(29), 4752 (2021). doi:10.1126/sciadv.abf4752

20. Sanz Leon, P., Knock, S., Woodman, M., Domide, L., Mersmann, J., McIntosh, A., Jirsa, V.: The virtual brain: a simulator of primate brain network dynamics. Frontiers in Neuroinformatics 7, 10 (2013). doi:10.3389/fninf.2013.00010

21. Cabral, J., Kringelbach, M.L., Deco, G.: Functional connectivity dynamically evolves on multiple time-scales over a static structural connectome: Models and mechanisms. NeuroImage 160, 84–96 (2017). doi:10.1016/j.neuroimage.2017.03.045

22. McKhann, G., Drachman, D., Folstein, M., Katzman, R., Price, D., Stadlan, E.M.: Clinical diagnosis of alzheimer’s disease: Report of the NINCDS-ADRDA work group under the auspices of department of health and human services task force on alzheimer’s disease. Neurology 34(7), 939–939 (1984). doi:10.1212/wnl.34.7.939

23. Glasser, M.F., Sotiropoulos, S.N., Wilson, J.A., Coalson, T.S., Fischl, B., Andersson, J.L., Xu, J., Jbabdi, S., Webster, M., Polimeni, J.R., Essen, D.C.V., Jenkinson, M.: The minimal preprocessing pipelines for the human connectome project. NeuroImage 80, 105–124 (2013). doi:10.1016/j.neuroimage.2013.04.127

24. Reuter, M., Schmansky, N.J., Rosas, H.D., Fischl, B.: Within-subject template estimation for unbiased longitudinal image analysis. NeuroImage 61(4), 1402–1418 (2012). doi:10.1016/j.neuroimage.2012.02.084

25. Smith, S.M., Jenkinson, M., Woolrich, M.W., Beckmann, C.F., Behrens, T.E.J., Johansen-Berg, H., Bannister, P.R., Luca, M.D., Drobnjak, I., Flitney, D.E., Niazy, R.K., Saunders, J., Vickers, J., Zhang, Y., Stefano, N.D., Brady, J.M., Matthews, P.M.: Advances in functional and structural MR image analysis and implementation as FSL. NeuroImage 23, 208–219 (2004). doi:10.1016/j.neuroimage.2004.07.051

26. Woolrich, M.W., Jbabdi, S., Patenaude, B., Chappell, M., Makni, S., Behrens, T., Beckmann, C., Jenkinson, M., Smith, S.M.: Bayesian analysis of neuroimaging data in FSL. NeuroImage 45(1), 173–186 (2009). doi:10.1016/j.neuroimage.2008.10.055

27. Jenkinson, M., Beckmann, C.F., Behrens, T.E.J., Woolrich, M.W., Smith, S.M.: FSL. NeuroImage 62(2), 782–790 (2012). doi:10.1016/j.neuroimage.2011.09.015

28. Glasser, M.F., Coalson, T.S., Robinson, E.C., Hacker, C.D., Harwell, J., Yacoub, E., Ugurbil, K., Andersson, J., Beckmann, C.F., Jenkinson, M., Smith, S.M., Essen, D.C.V.: A multi-modal parcellation of human cerebral cortex. Nature 536(7615), 171–178 (2016). doi:10.1038/nature18933

29. Robinson, E.C., Jbabdi, S., Glasser, M.F., Andersson, J., Burgess, G.C., Harms, M.P., Smith, S.M., Essen, D.C.V., Jenkinson, M.: MSM: A new flexible framework for multimodal surface matching. NeuroImage 100, 414–426 (2014). doi:10.1016/j.neuroimage.2014.05.069

30. Veraart, J., Novikov, D.S., Christiaens, D., Ades-aron, B., Sijbers, J., Fieremans, E.: Denoising of diffusion MRI using random matrix theory. NeuroImage 142, 394–406 (2016). doi:10.1016/j.neuroimage.2016.08.016

31. Tournier, J.-D., Calamante, F., Connelly, A.: Determination of the appropriate value and number of gradient directions for high-angular-resolution diffusion-weighted imaging. NMR in Biomedicine 26(12), 1775–1786 (2013). doi:10.1002/nbm.3017

32. Tournier, J.-D., Calamante, F., Connelly, A.: Robust determination of the fibre orientation distribution in diffusion MRI: Non-negativity constrained super-resolved spherical deconvolution. NeuroImage 35(4), 1459–1472 (2007). doi:10.1016/j.neuroimage.2007.02.016

33. Tournier, J.D., Calamante, F., Connelly, A.: Improved probabilistic streamlines tractography by 2nd order integration over fibre orientation distributions. In: Proceedings of the International Society for Magnetic Resonance in Medicine, vol. 18, p. 1670 (2010)

34. Smith, R.E., Tournier, J.-D., Calamante, F., Connelly, A.: SIFT2: Enabling dense quantitative assessment of brain white matter connectivity using streamlines tractography. NeuroImage 119, 338–351 (2015). doi:10.1016/j.neuroimage.2015.06.092

35. Smith, R.E., Tournier, J.-D., Calamante, F., Connelly, A.: Anatomically-constrained tractography: Improved diffusion MRI streamlines tractography through effective use of anatomical information. NeuroImage 62(3), 1924–1938 (2012). doi:10.1016/j.neuroimage.2012.06.005

36. Wong, K.-F., Wang, X.-J.: A recurrent network mechanism of time integration in perceptual decisions. Journal of Neuroscience 26(4), 1314–1328 (2006). doi:10.1523/jneurosci.3733-05.2006

37. Stephan, K.E., Kasper, L., Harrison, L.M., Daunizeau, J., den Ouden, H.E.M., Breakspear, M., Friston, K.J.: Nonlinear dynamic causal models for fMRI. NeuroImage 42(2), 649–662 (2008). doi:10.1016/j.neuroimage.2008.04.262

38. Fu, H., Possenti, A., Freer, R., Nakano, Y., Villegas, N.C.H., Tang, M., Cauhy, P.V.M., Lassus, B.A., Chen, S., Fowler, S.L., Figueroa, H.Y., Huey, E.D., Johnson, G.V.W., Vendruscolo, M., Duff, K.E.: A tau homeostasis signature is linked with the cellular and regional vulnerability of excitatory neurons to tau pathology. Nature Neuroscience 22(1), 47–56 (2018). doi:10.1038/s41593-018-0298-7

39. Zott, B., Simon, M.M., Hong, W., Unger, F., Chen-Engerer, H.-J., Frosch, M.P., Sakmann, B., Walsh, D.M., Konnerth, A.: A vicious cycle of β amyloid–dependent neuronal hyperactivation. Science 365(6453), 559–565 (2019). doi:10.1126/science.aay0198

40. McInnes, J., Wierda, K., Snellinx, A., Bounti, L., Wang, Y.-C., Stancu, I.-C., Apóstolo, N., Gevaert, K., Dewachter, I., Spires-Jones, T.L., Strooper, B.D., Wit, J.D., Zhou, L., Verstreken, P.: Synaptogyrin-3 mediates presynaptic dysfunction induced by tau. Neuron 97(4), 823–8358 (2018). doi:10.1016/j.neuron.2018.01.022

41. Vasconcelos, B., Stancu, I.-C., Buist, A., Bird, M., Wang, P., Vanoosthuyse, A., Kolen, K.V., Verheyen, A., Kienlen-Campard, P., Octave, J.-N., Baatsen, P., Moechars, D., Dewachter, I.: Heterotypic seeding of tau fibrillization by pre-aggregated abeta provides potent seeds for prion-like seeding and propagation of tau-pathology in vivo. Acta Neuropathologica 131(4), 549–569 (2016). doi:10.1007/s00401-015-1525-x

42. Griner, S.L., Seidler, P., Bowler, J., Murray, K.A., Yang, T.P., Sahay, S., Sawaya, M.R., Cascio, D., Rodriguez, J.A., Philipp, S., Sosna, J., Glabe, C.G., Gonen, T., Eisenberg, D.S.: Structure-based inhibitors of amyloid beta core suggest a common interface with tau. eLife 8, 46924 (2019). doi:10.7554/eLife.46924

43. Ünal Sakoğlu Pearlson, G.D., Kiehl, K.A., Wang, Y.M., Michael, A.M., Calhoun, V.D.: A method for evaluating dynamic functional network connectivity and task-modulation: application to schizophrenia. Magnetic Resonance Materials in Physics, Biology and Medicine 23(5-6), 351–366 (2010). doi:10.1007/s10334-010-0197-8

44. Preti, M.G., Bolton, T.A., Ville, D.V.D.: The dynamic functional connectome: State-of-the-art and perspectives. NeuroImage 160, 41–54 (2017). doi:10.1016/j.neuroimage.2016.12.061

45. Faul, F., Erdfelder, E., Buchner, A., Lang, A.-G.: Statistical power analyses using g∗power 3.1: Tests for correlation and regression analyses. Behavior Research Methods 41(4), 1149–1160 (2009). doi:10.3758/brm.41.4.1149

46. Jansen, B.H., Rit, V.G.: Electroencephalogram and visual evoked potential generation in a mathematical model of coupled cortical columns. Biological Cybernetics 73(4), 357–366 (1995). doi:10.1007/bf00199471

47. Riley, K.P., Snowdon, D.A., Markesbery, W.R.: Alzheimer’s neurofibrillary pathology and the spectrum of cognitive function: Findings from the nun study. Annals of Neurology 51(5), 567–577 (2002). doi:10.1002/ana.10161

48. Wang, P., Kong, R., Kong, X., Liégeois, R., Orban, C., Deco, G., van den Heuvel, M.P., Yeo, B.T.T.: Inversion of a large-scale circuit model reveals a cortical hierarchy in the dynamic resting human brain. Science Advances 5(1), 7854 (2019). doi:10.1126/sciadv.aat7854

49. Chaudhuri, R., Knoblauch, K., Gariel, M.-A., Kennedy, H., Wang, X.-J.: A large-scale circuit mechanism for hierarchical dynamical processing in the primate cortex. Neuron 88(2), 419–431 (2015). doi:10.1016/j.neuron.2015.09.008

50. Marquié, M., Verwer, E.E., Meltzer, A.C., Kim, S.J.W., Agüero, C., Gonzalez, J., Makaretz, S.J., Chong, M.S.T., Ramanan, P., Amaral, A.C., Normandin, M.D., Vanderburg, C.R., Gomperts, S.N., Johnson, K.A., Frosch, M.P., Gómez-Isla, T.: Lessons learned about [f-18]-AV-1451 off-target binding from an autopsy-confirmed parkinson’s case. Acta Neuropathologica Communications 5(1) (2017). doi:10.1186/s40478-017-0482-0

51. Ikonomovic, M.D., Abrahamson, E.E., Price, J.C., Mathis, C.A., Klunk, W.E.: [f-18]AV-1451 positron emission tomography retention in choroid plexus: More than “off-target” binding. Annals of Neurology 80(2), 307–308 (2016). doi:10.1002/ana.24706

52. Fruchterman, T.M.J., Reingold, E.M.: Graph drawing by force-directed placement. Software: Practice and Experience 21(11), 1129–1164 (1991). doi:10.1002/spe.4380211102

